# Inclusion, reporting and analysis of demographic variables in chronobiology and sleep research

**DOI:** 10.1101/2023.03.28.534522

**Authors:** Selma Tir, Rhiannon White, Manuel Spitschan

**Affiliations:** Department of Experimental Psychology, University of Oxford; Sleep and Circadian Neuroscience Institute, Nuffield Department of Clinical Neurosciences, University of Oxford; Warwick Medical School, University of Warwick, United Kingdom; Department of Health and Sport Sciences, TUM School of Medicine and Health, Technical University of Munich, Munich, Germany; Translational Sensory & Circadian Neuroscience, Max Planck Institute for Biological Cybernetics, Tübingen, Germany; TUM Institute of Advanced Study (TUM-IAS), Technical University of Munich, Garching, Germany

**Keywords:** demographics, ethnicity, sex, research participants, reporting, publishing, meta-science

## Abstract

Many aspects of sleep and circadian physiology appear to be sensitive to participant-level characteristics. While recent research robustly highlights the importance of considering participant-level demographic information, it is not clear to what extent this information is available within the large body of existing literature. This article investigates study sample characteristics within the published sleep and chronobiology research over the past 40 years. 6,777 articles were identified and a random sample of 20% was included. The reporting of sample size, age, sex, gender, ethnicity, level of education, socio-economic status, and profession of the study population was scored, and any reported aggregate summary statistics for these variables were recorded. We found that while >90% of studies reported age or sex, all other variables were reported in <25% of cases. Reporting quality was highly variable, indicating an opportunity to standardize reporting guidelines for participant-level characteristics to facilitate meta analyses.

**Summary:** In this article, we address the question of how representative, diverse and inclusive are published articles in sleep and chronobiology research. We analyzed a sample of >1300 articles published in sleep and chronobiology journals between 1979 and 2019 for its inclusion, reporting and analysis of study population characteristics, including age, sex, gender, race/ethnicity, level of education, socio-economic status, and profession. We found that while >90% of studies reported age or sex, all other variables were reported in <25% of cases, with the frequency of reporting changing over time. We identify opportunities for improving the reporting of demographic variables.

**Research Agenda:** Future research needs to: 1. Establish schemas for reporting demographic variables in a harmonized way across geographical and cultural contexts; 2. Identify gaps in the sleep and chronobiology literature with respect to understudied populations; 3. Understand the extent to which research practices allow for the inclusion of diverse populations in all stages of the research cycle, and how this can exacerbate health inequities.

**Practice Points:**

1. Published studies on circadian and sleep physiology should be carefully examined.
2. Reporting of demographic variables should be done deliberately and systematically.
3. Inclusion and diversity of different populations across the field needs to be ensured.

## Introduction

Sleep and circadian rhythms are essential physiological and behavioral processes that have been shown to vary significantly across individuals (1–7). People can differ in their sleep patterns, including the amount, timing, and quality of sleep they require, as well as circadian rhythms, such as chronotype and period. Some of these differences have been systematically related to demographic variables, most notably age (8–14), sex (15–19) and ethnicity (20–23), demonstrating the need to consider sleep and circadian physiology at a participant-level. For example, children and adolescents have different sleep patterns compared to adults, with the *need for sleep* generally declining with age and shifting towards a morning chronotype (**yoon_2001?**). On average, women have a shorter circadian period (24), may be more likely to experience sleep disturbances and insomnia (25), but are less prone to poorer sleep with aging compared to men (15). The impact of gender-related factors such as menstrual cycles, pregnancy, and menopause can also inform the development of sex-specific interventions and improve the diagnosis and treatment of sleep disorders in both sexes (26). Studies have shown that sleep problems and circadian rhythm disruptions are more common among certain racial/ethnic groups, such as African Americans and Hispanics/Latinos, compared to non-Hispanic Whites. These disparities have been observed through differences in circadian period, chronotype, phase shifting response and sleep duration (27).

Zooming out from the level of individual studies, there are significant inequities in sleep health depending on education level, profession and socio-economic status (28–33). These factors can influence an individual’s access to resources and better living conditions, which can in turn affect their sleep and circadian rhythms (34). For example, it is suggested that individuals with lower SES are more likely to experience sleep disturbances and circadian rhythm disruptions compared to those with higher SES (35). This may be due to a number of factors, such as stress, work demands and access to healthcare. They may also live in neighborhoods with more noise pollution or exposure to artificial light. Similarly, different professions or employment statuses can have varying work schedules, demands, and stress levels. Individuals who work night shifts, irregular schedules, or long hours may experience circadian misalignment and have difficulty sleeping, which in turn leads to adverse health effects. Geographical location may also be a relevant factor for sleep and circadian rhythms research since it can influence environmental factors such as natural light exposure, temperature, altitude, noise levels, and air pollution. In countries with a hotter climate, it is common to take a midday nap or *siesta*, which can affect sleep patterns and alter the timing of the body’s internal clock. Some individual aspects of sleep and circadian physiology may thus be linked to genetic predispositions or influenced by cultural, environmental and societal factors. Population-based studies may be key to offer personalized solutions and treatments for sleep and circadian disruption. As compromised sleep has many knock-on effects, including negative effects on cardiovascular, metabolic, neurobehavioral and cognitive function, it is imperative to understand how demographic variables influence sleep and circadian rhythms.

The extent to which a scientific field’s findings are generalizable is a function of the representativeness of a given study sample, and the degree to which findings differ between, e.g., different sexes and ethnicities, also referred to as external validity. It is suggested that research practices have historically excluded diverse populations in all stages of the research cycle, including recruitment, retention, data collection, analysis, and dissemination of findings. In the domain of sex, a recent study reviewed the reporting and analysis of sex in biological sciences research (36). The authors found that while sex inclusion has significantly increased over the past 10 years (37), sex-based analysis has not improved despite recent policies and funder mandates (38). This call for inclusion outlined several guidelines, including involving diverse communities in the design and implementation of research studies, using culturally sensitive recruitment and retention strategies, and reporting demographic data in a transparent and standardized manner. Indeed, in 2016 the United States’ National Institutes of Health (NIH) issued a notice requiring grant holders to factor sex into the design, analysis and reporting of vertebrate and human studies, or to provide substantial justification for the study of a single sex (39). The term “gender data gap” has recently been introduced, demonstrating that women have historically been excluded from biomedical research (40). Similarly, the lack of diversity in biomedical and clinical research, and understudy of minorities in the presence of existing health inequities exacerbates these inequities (41). A recent review of contemporary dementia research reported a lack of demographic, racial, and geographic diversity (42). Similarly, it has been reported that the study populations of clinical trials of 53 drugs approved by the US Food and Drug Administration (FDA) in 2020 consisted of 75% White (43). Together, these findings outline the need for representativeness in study populations in order to develop effective and equitable tailored interventions to promote healthy sleep and circadian rhythms for all.

While research findings converge on participant-level demographic characteristics affecting outcomes, it is not clear to what extent this information is available in the large body of historic literature, nor to what extent it is even reported. Here, we address the question of participant-level demographic characteristics (age, sex, gender, ethnicity, level of education, socio-economic status, and profession of the study population) and reporting thereof in chronobiology and sleep research over the past fourty years. We extracted the study sample characteristics in a total of 1355 randomly sampled publications across the eight top (ranked by the Journal Impact Factor) chronobiology and sleep research journals, and subjected them to a comprehensive analysis in terms of the inclusion, reporting and analysis of demographic variables.

## Methods

### Procedure

Journal articles published between 1979 and 2019 in the top eight sleep and chronobiology journals were considered. For practical reasons, a temporal resolution of 2 years was considered sufficient to determine any effects changing over time, and the number of screened articles was reduced by only analyzing those published in odd years. The list of possible target journals was based on a previously established list of journals implementing a hybrid strategy by consulting the Web of Science Master Journal List, domain-relevant expertise in sleep and chronobiology and consulting with a senior researcher with >25 years of experience in the field (44). From this previously derived list, we selected eight journals based on their five-year Impact Factor, and included *Journal of Pineal Research* (ISSN: 0742-3098 / 1600-079X; 2018 5-year IF: 12.197), *Sleep* (0161-8105 / 1550-9109; 5.588), *Journal of Sleep Research* (0962-1105 / 1365-2869; 3.951), *Sleep Medicine* (1389-9457 / 1878-5506; 3.934), *Journal of Clinical Sleep Medicine* (1550-9389 / 1550-9397; 3.855), *Journal of Biological Rhythms* (0748-7304 / 1552-4531; 3.349), Behavioral Sleep Medicine (1540-2002 / 1540-2010; 3.162), and *Chronobiology International* (0742-0528 / 1525-6073; 2.998). While *Sleep Medicine Reviews* also features in the list of journals, we did not include it as it primarily publishes reviews.

### Article inclusion

6,777 articles were identified through a MEDLINE search and filtering by journal and odd years. A random sample of 20% was initially selected for screening. Inclusion requirements included conducting original research in the English language, reporting human data, and recruiting volunteers. As such, animal studies, bibliographies, case reports, comments, conference proceedings, editorials, guidelines, letters, retracted publications, reviews, errata and corrigenda were excluded.

### Review and article extraction

All included articles were reviewed for eligibility and coded by RW. The reporting of sample size, age, sex, gender, race/ethnicity, level of education, socio-economic status, and profession of the study population was scored binarily (0 = not reported, 1 = reported), and any reported aggregate summary statistics for these variables were recorded (e.g., mean, median, etc.). Sample size referred to the total number of participants for a given study. If an article reported multiple studies, then it was analyzed for each of its individual studies. Age was analyzed when recorded in days, weeks or years. Sex referred to biological sex, while gender referred to the social construction of sex. When sex and gender were used interchangeably and didn’t refer to personal identification, it was scored as biological sex. Since the language and system describing ethnicity, level of education, socio-economic status and profession differ between countries and individuals, all reporting was taken into account as long as there was a clear indication of what the variable represented. For example, socio-economic status included categories of income, seniority within a company or type of labor, and “occupation” and “employment status” were recorded as profession. Additionally, the non-demographic variables, funding source, geographical location and clinical focus of the article, were examined, as well as whether data were analyzed by including any of the demographic variables as covariates. Data were coded in an Excel Spreadsheet and analyzed in R Studio (version 4.2.2).

### Pre-registration

We pre-registered our protocol (specified using the PRISMA-P template (45,46)) on the Open Science Framework (https://osf.io/cu3we/). For thorough details on the screening and analysis methods, see the protocol.

### Materials, data and code availability

All data underlying this manuscript are available on a public GitHub repository (https://github.com/MPI-tSCN/sleep_circadian_demographics_data). The article was written in R (47) using RMarkdown and papaja (48), employing a series of additional R packages (49–60) and is fully reproducible.

## Results

### Number of analyzed articles

Out of 1355 identified and pre-screened articles, we included and extracted data from 1152 (85%), following our inclusion requirements. The distribution of years in which the articles were published is non-uniform and the proportion of included and extracted data is greater for more recent articles (Fig. 1). In addition, the representation of journals in the final list of articles is also non-uniform, as the included journals were not all available from the start date of our data collection (1979 onwards).

**Figure 1.**
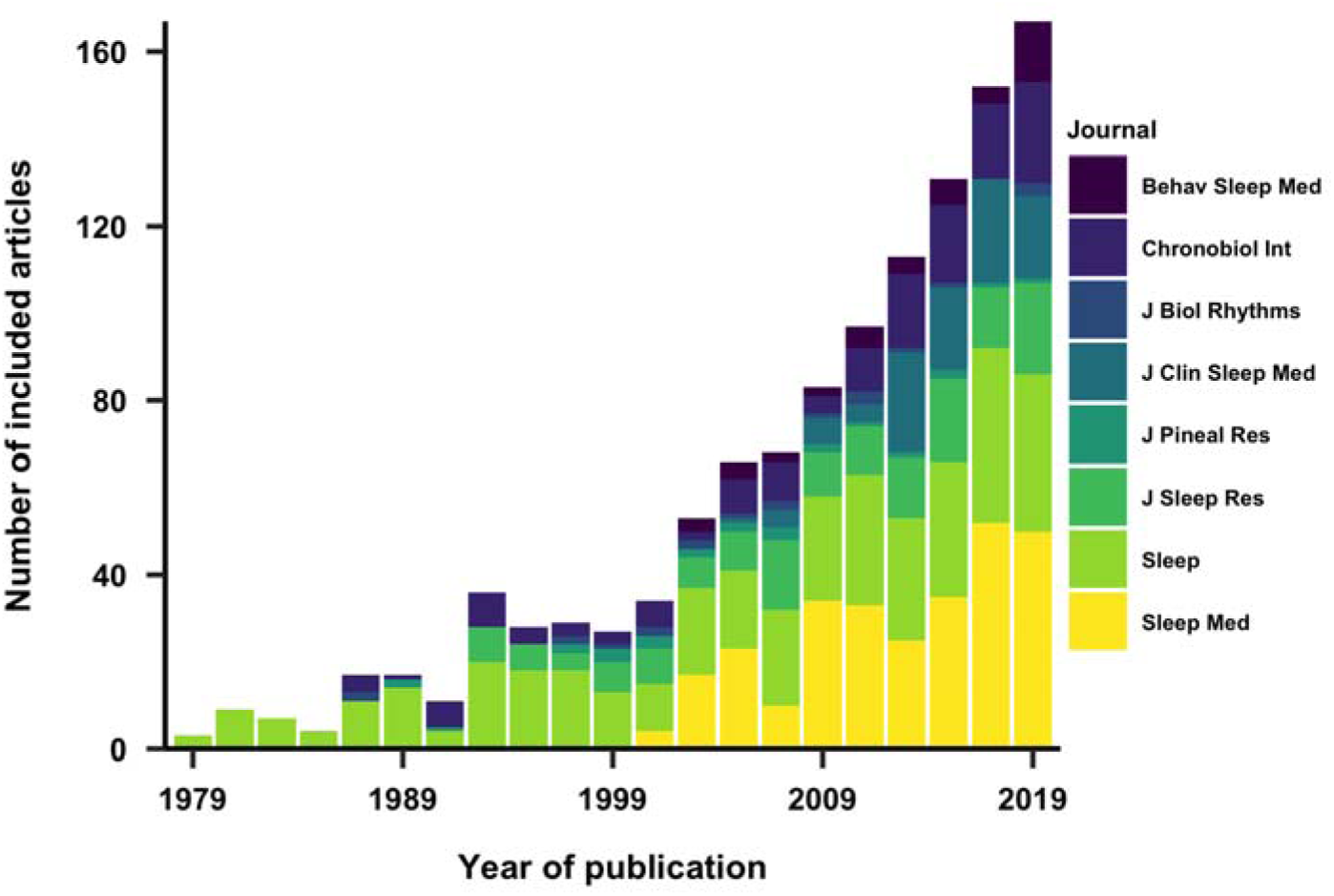
Included and analyzed articles by year and journal of publication. Recent articles are more widely represented, reflecting an overall increase in scientific output.

We also investigated reasons for exclusion among the articles that we did not include or extract data from. These are given in Figure 2. Reasons for exclusion vary somewhat between different years, with articles not reporting original research being excluded at the highest rate on average.

**Figure 2.**
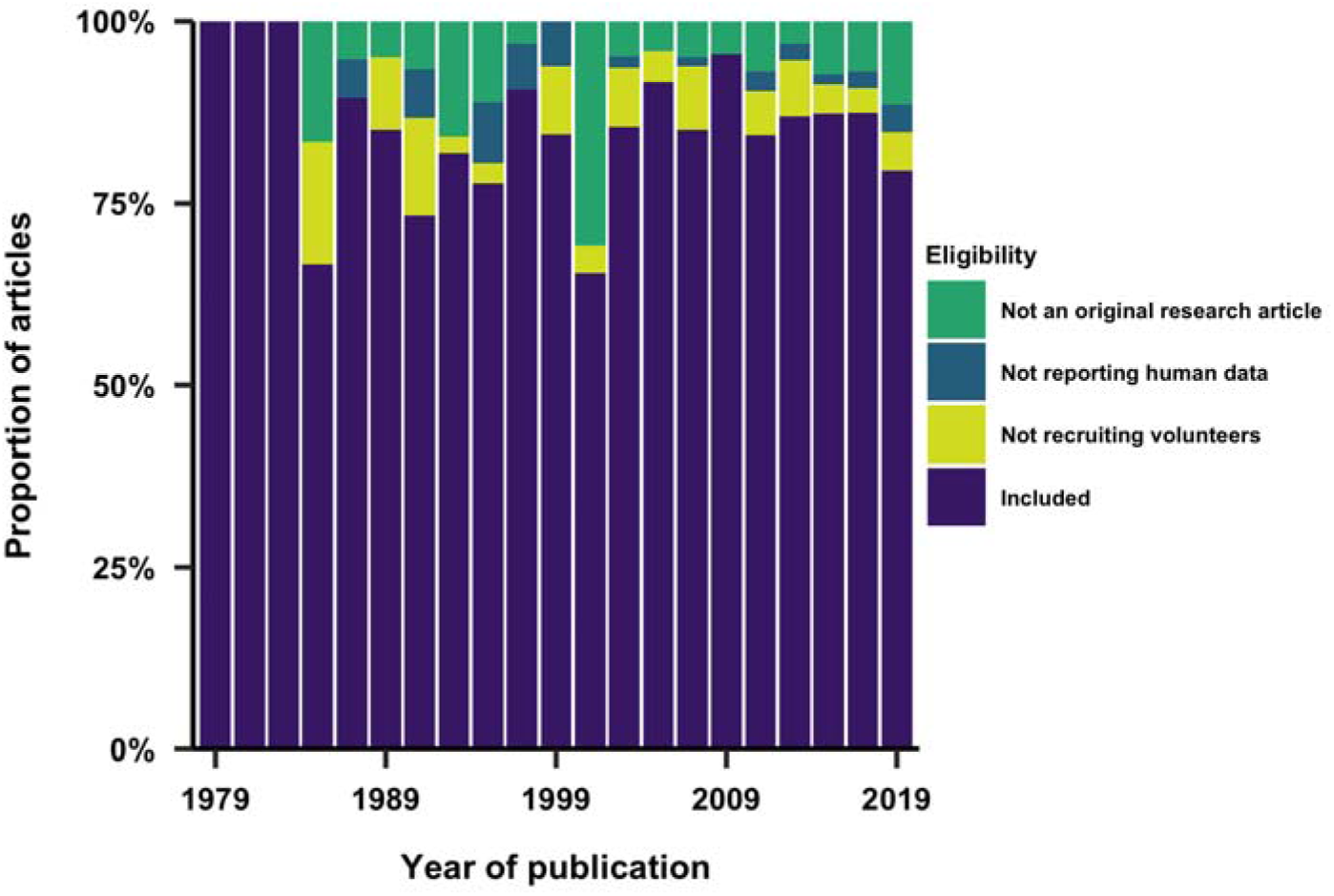
Percentage of included and excluded articles by year of publication, with reasons for exclusion.

### Funding

We examined the reporting of funding sources in included articles. Funding sources were reported by 62% of studies, while funding number was also reported in 69% of these cases (Fig. 3). Overall, funding by the United States’ National Institutes of Health (NIH) represented 19% of the reported funding agencies. 92% of the studies funded by the NIH also reported funding number. The second most represented funding agencies were the Australian National Health and Medical Research Council (NHMRC) and the Canadian Institutes of Health Research (CIHR).

**Figure 3.**
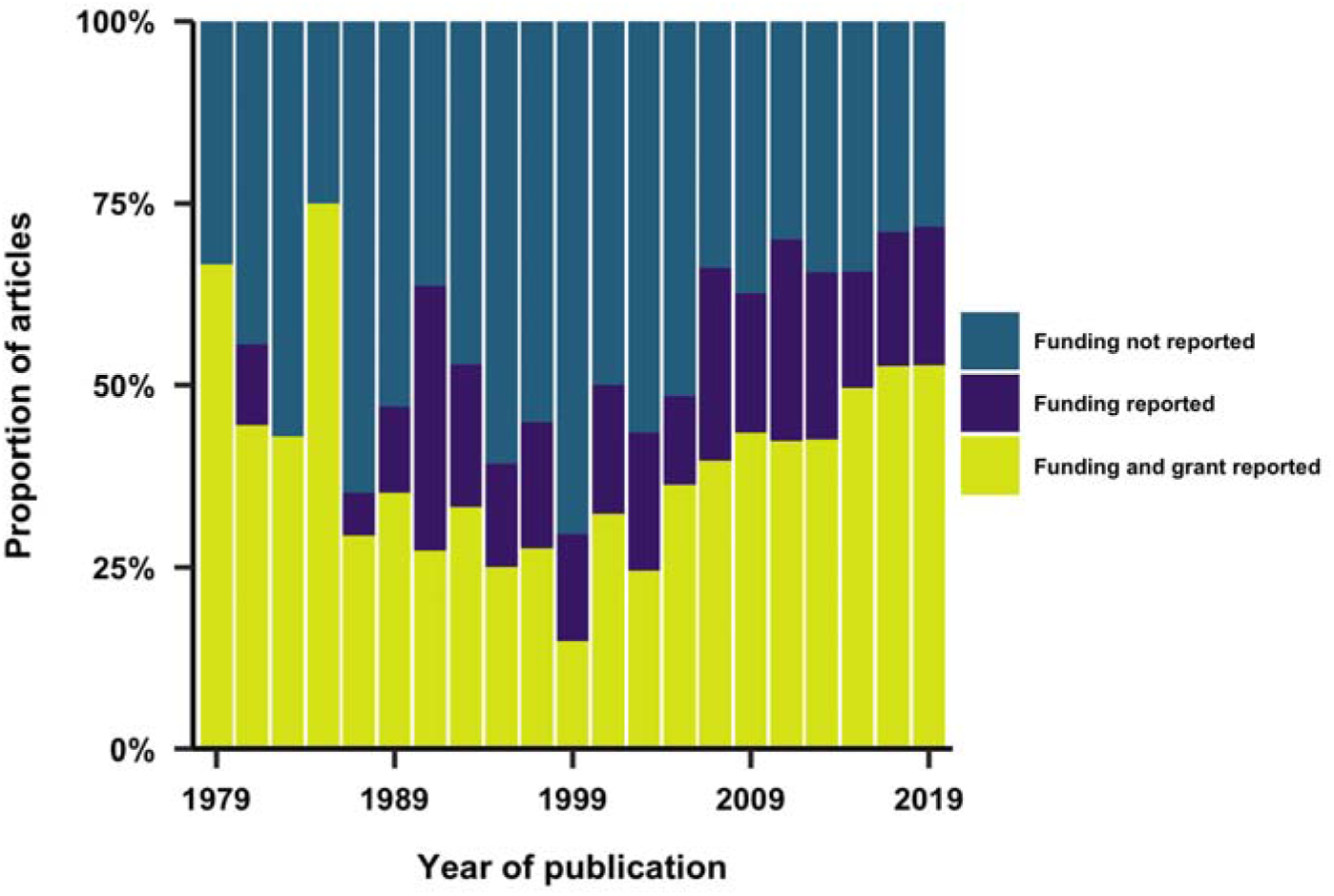
Reporting of funding across years.

### Geographical location

93% of articles were conducted in a single country. The geographical location of the study was explicitly reported in 57% of studies. The country of study was inferred for the rest of the sampled articles. Inference was primarily based on the first author’s affiliation as it typically indicates the institution where the research was conducted at the time of the study. We also speculated that if the study population was not that of the institution’s geographic location, then it would be clearly stated in the article. Overall, 53 countries were represented. Figure 4 shows the distribution of study location across time with the eight most represented countries. Across all years, the US is the most represented country and only three studies were primarily conducted in Africa (Nigeria, Senegal and Tunisia).

**Figure 4.**
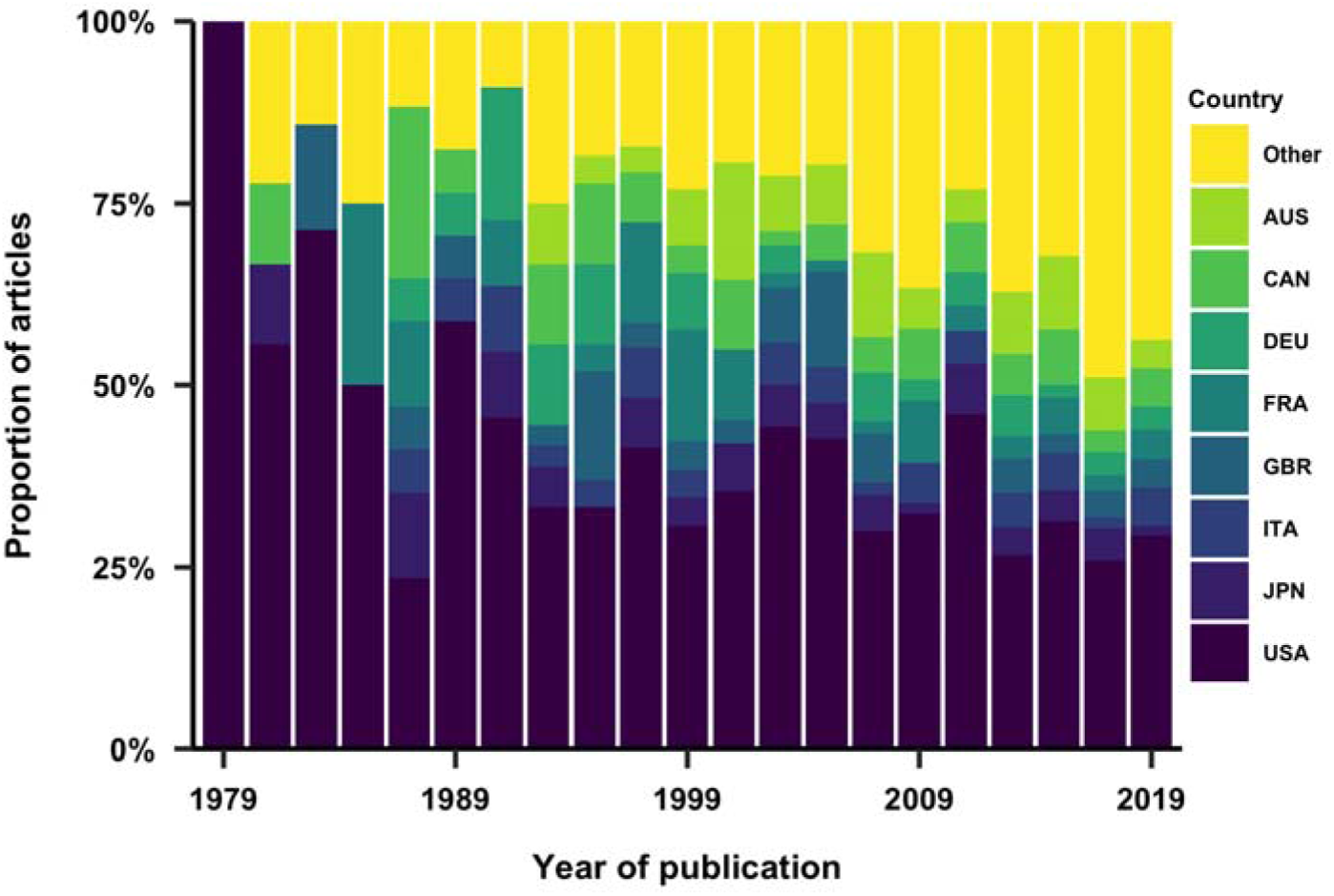
Geographical location of the studies. The eight most represented countries across the entire dataset are individually shown.

### Sample size

We examined whether sample size was reported in studies, and the numbers recorded. Overall, sample size was reported in 92% of studies. We furthermore examined the distribution of sample sizes as a function of the publication year of the article (Fig. 5), showing a wider distribution of sample sizes in more recent articles.

**Figure 5.**
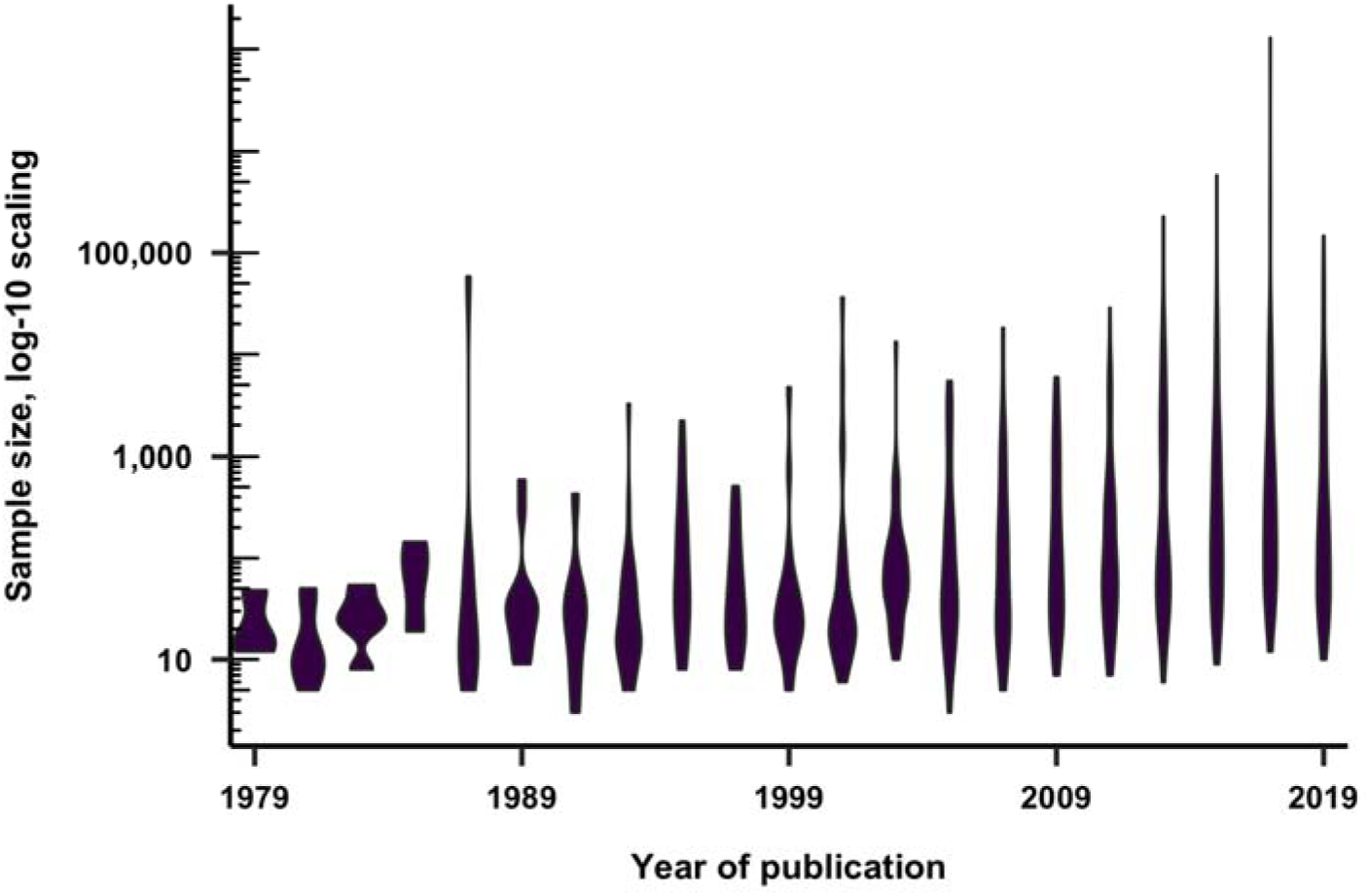
Sample size of the recruited volunteers as a function of publication year. Numbers are computed on a log-10 scale.

### Age

93% of articles reported a variable describing age, such as the mean, standard deviation, median, minimum, maximum and interquartile range. Overall, the median and interquartile range were the least reported variables for age (Fig. 6). The minimum and maximum age were the most employed variables in 1979, while their reporting decreased throughout the years compared to other variables. On another hand, the standard deviation of the mean was increasingly reported throughout the years. Trends for specific journals were also observed, such as the substantial use of the min/max variables in *Journal of Pineal Research*, and the lack of in *Journal of Clinical Sleep Medicine*.

**Figure 6.**
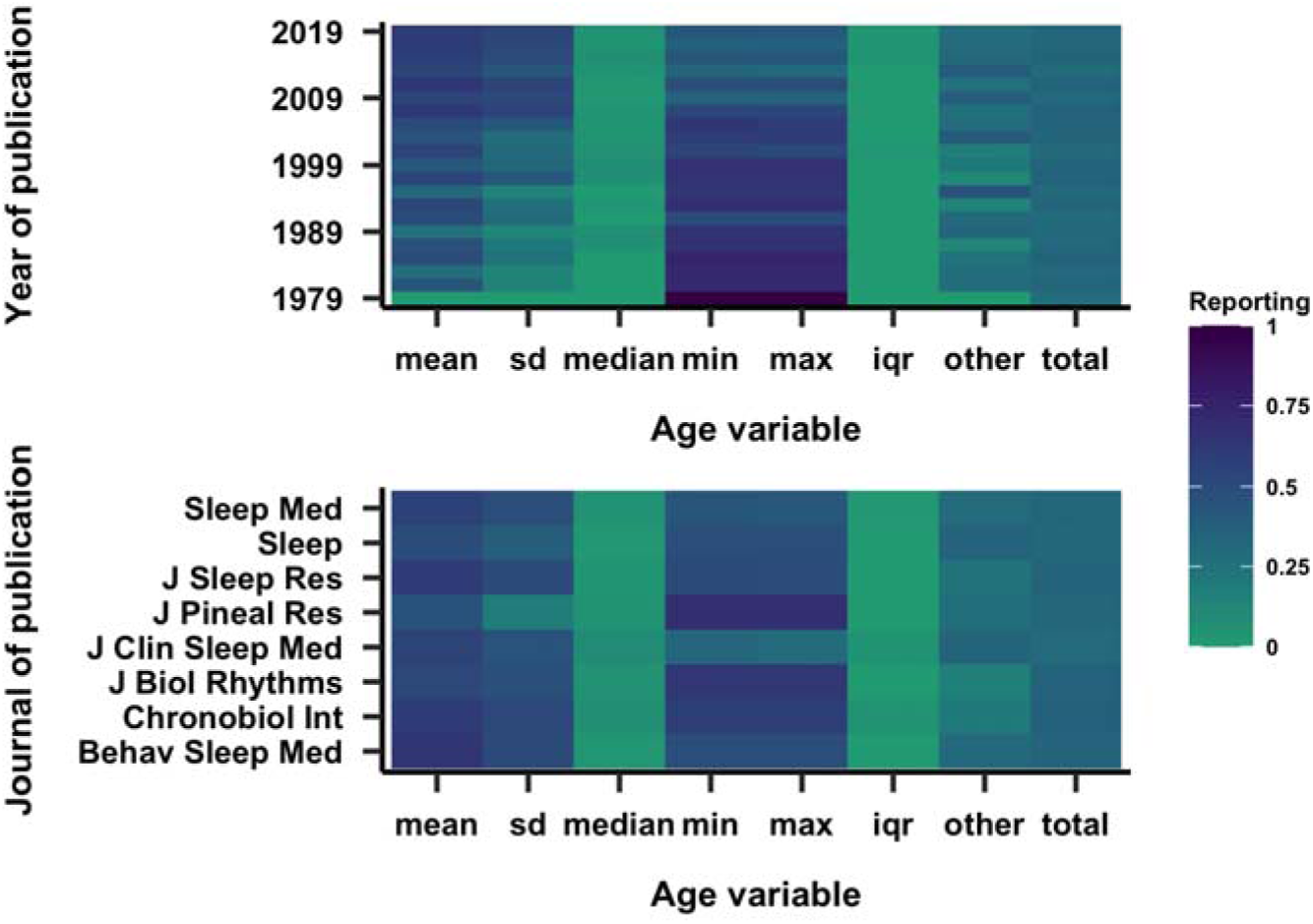
Reporting of various age variables by year (top) and journal (bottom) of publication. Darker shades imply a higher correlation. SD = standard deviation of the mean, IQR = interquartile range.

Overall, the average mean age of the study populations was 39 years old. We examined the extent to which the mean age differed across studies as a function of publication year (Fig. 7), and found that the mean age is much more widely varied in later years, likely reflecting the extent of considering study samples that are more diversified in age. It is interesting to note that while the range of ages significantly expands throughout the years, the mean age remains somewhat constant, centered around the 40 year old population.

**Figure 7.**
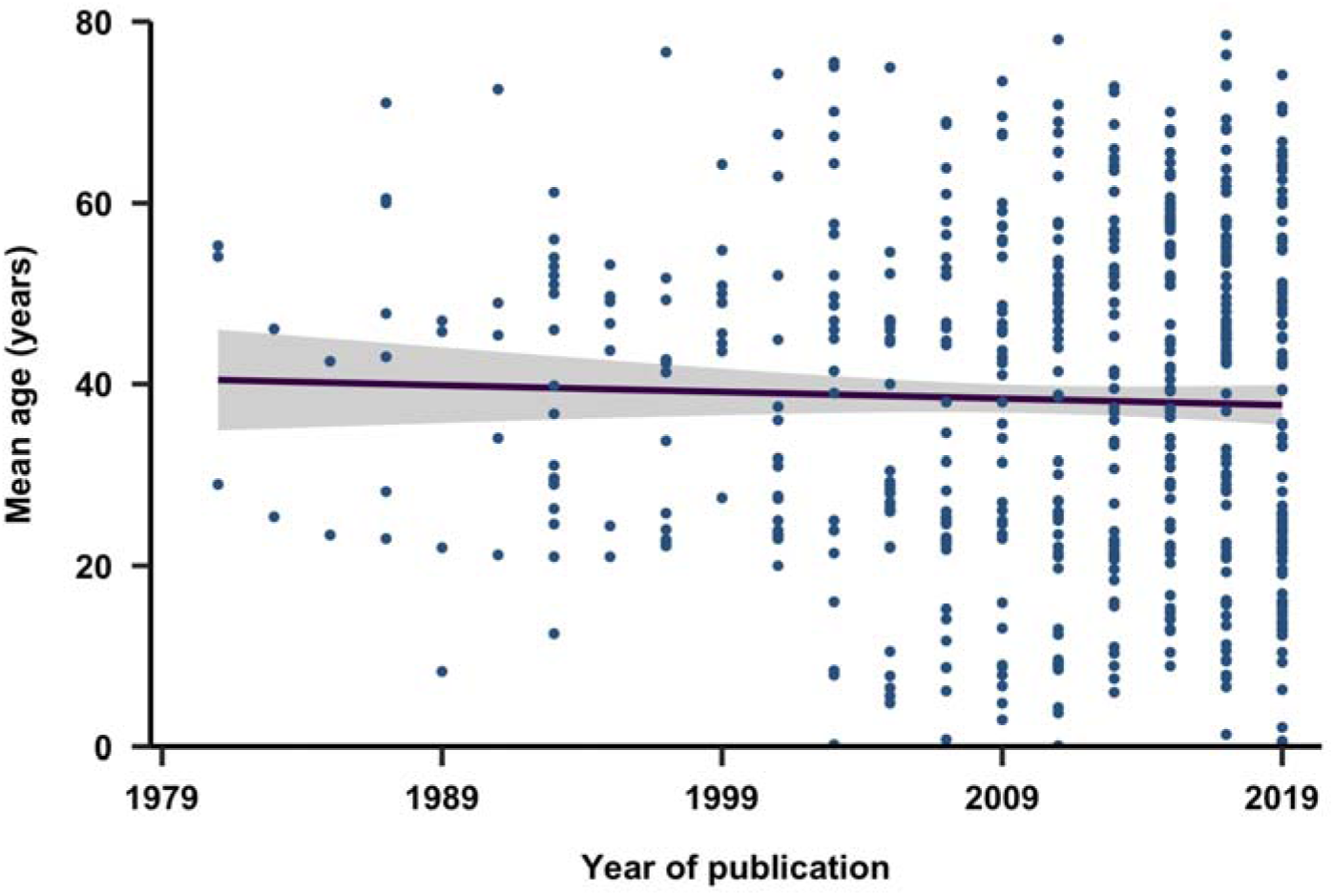
Development of mean age in included studies as a function of publication year. Fit shown is a linear fit (±95%CI).

### Sex & Gender

Sex was reported in 89% of the studies. In Figure 8, we show the proportion of studies that recruited male subjects, female subjects, both sexes, or did not specify the sex of the participants. Overall, 13% of the studies reporting sex only recruited male participants, while 10% only employed females. Out of the studies focusing on a single sex, 1% of the male studies focused on a sex dependent feature, while 2% of the female studies did. 4% of studies reported age by sex.

**Figure 8.**
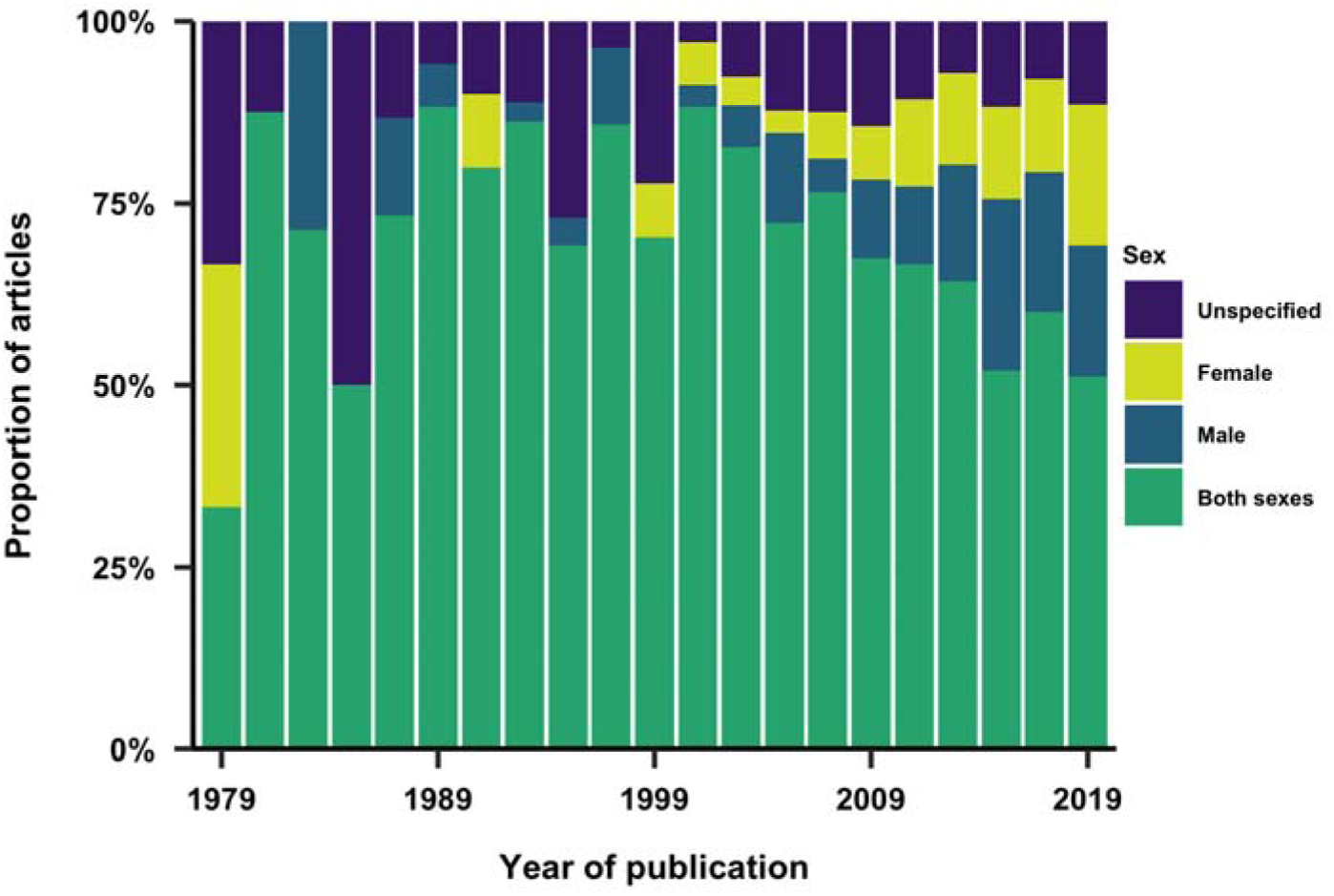
Sex inclusion by year. Proportion of studies that recruited male subjects, female subjects, both sexes, or did not specify the sex of the participants.

Differentiating between sex and gender may allow for a more comprehensive understanding of the potential differences in health outcomes between males and females, and the impact of social and cultural factors related to gender identity. Sex refers to the biological differences between males and females, while gender refers to the social and cultural roles, behaviors, and identities that are associated with being male or female. Incorporating sex and gender considerations can help ensure that treatments are safe and effective for all individuals, regardless of their sex or gender identity. Here, we found that gender was reported in only one article, with participants being described as male, female or transgender. This finding may suggest a lack of inclusion for gender identity and the transgender population.

### Ethnicity, education, profession and socio-economic status

We examined the reporting of additional demographic variables, including race/ethnicity, education, profession and socio-economic status (SES). Profession refers to an individual’s occupation or employment status, while SES refers to an individual’s or a family’s social and economic status. Education level, profession and SES can influence access to resources, opportunities, and living conditions. Understanding the impact of these variables can help identify groups that may be at higher risk for sleep-related conditions, and inform interventions to improve sleep and circadian rhythms in these populations. It can also help to address health disparities related to sleep and circadian rhythms by addressing underlying cultural, educational, socioeconomic and work-related factors that may be contributing to these disparities.

We found that these demographic variables were reported in 12% of studies for education, 15% for ethnicity, 4% for socio-economic status and 2% for profession. Figure 9 shows the distribution of this reporting across the years. Qualitatively, there is a clear increase in the reporting of additional demographic variables over time, with ethnicity being the most reported demographic variable across articles in more recent years.

**Figure 9.**
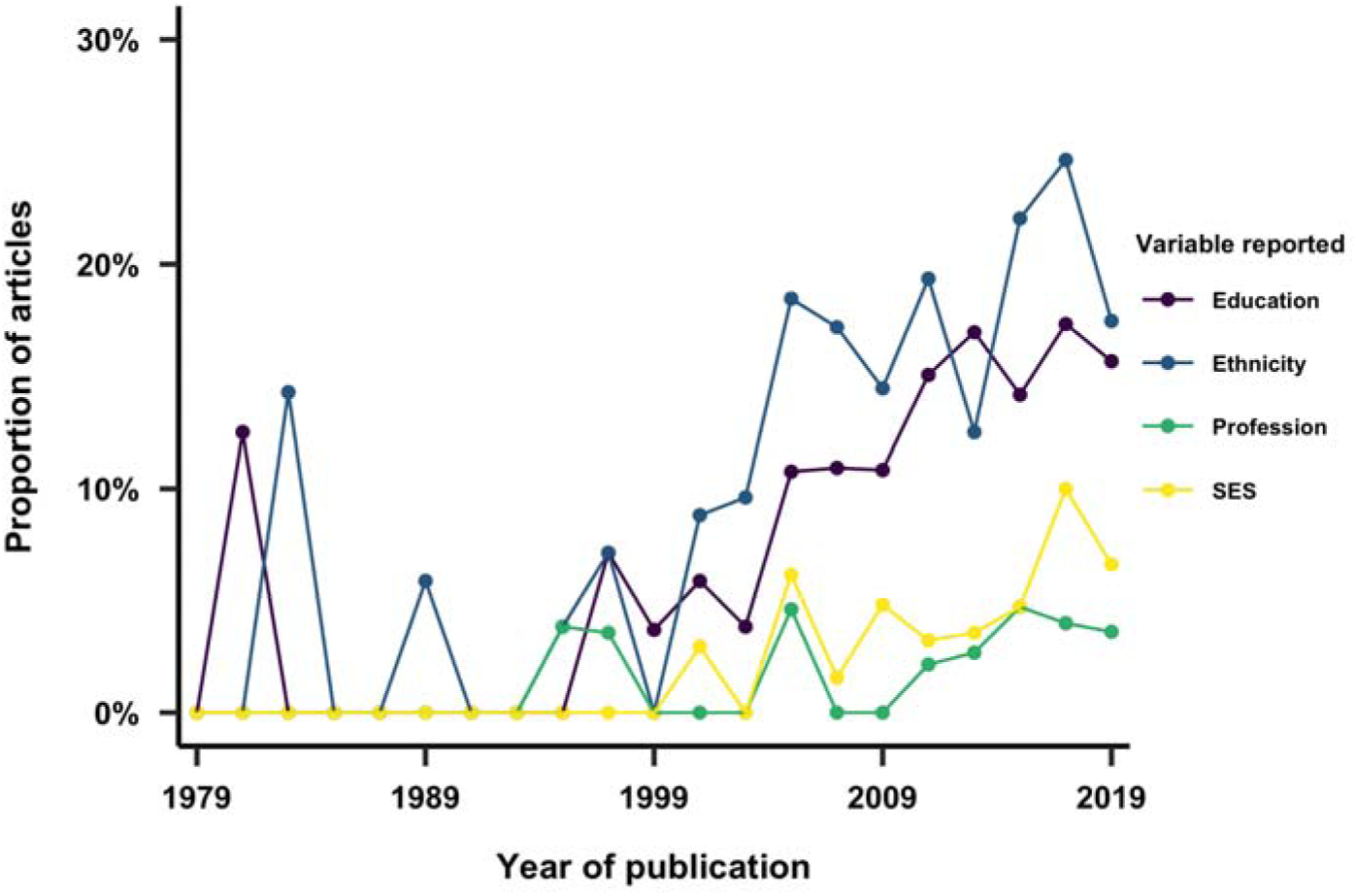
Reporting of education, ethnicity, profession and socio-economic status by year of publication.

We furthermore examined the number of categories included for each of these demographic variables. Figure 10 shows the number of categories reported for each variable among the articles that included it. On overage, we find that the coding scheme for each of these comprised of three to four categories. Education was reported as a range of years of study, degree level, or a subjective measure such as “low, medium, high.” The mean number of years of education was reported in 34% of articles, and was 14 years of study overall. Participants could choose multiple categories for ethnicity in 6% of the studies reporting this variable. The most common categories for ethnicity were “White/Caucasian,” “Black/African American” and “other,” and were mentioned in 89%, 60% and 53% of studies respectively. The most common category for profession was “unemployed,” mentioned in 39% of studies, along with a variety of specific jobs, such as attorney, farmer and astronaut. In 57% of studies reporting socio-economic status, the categories of SES were based on a range of income. The rest of the articles reported SES in subjective measures, such as “high, medium, low.”

**Figure 10.**
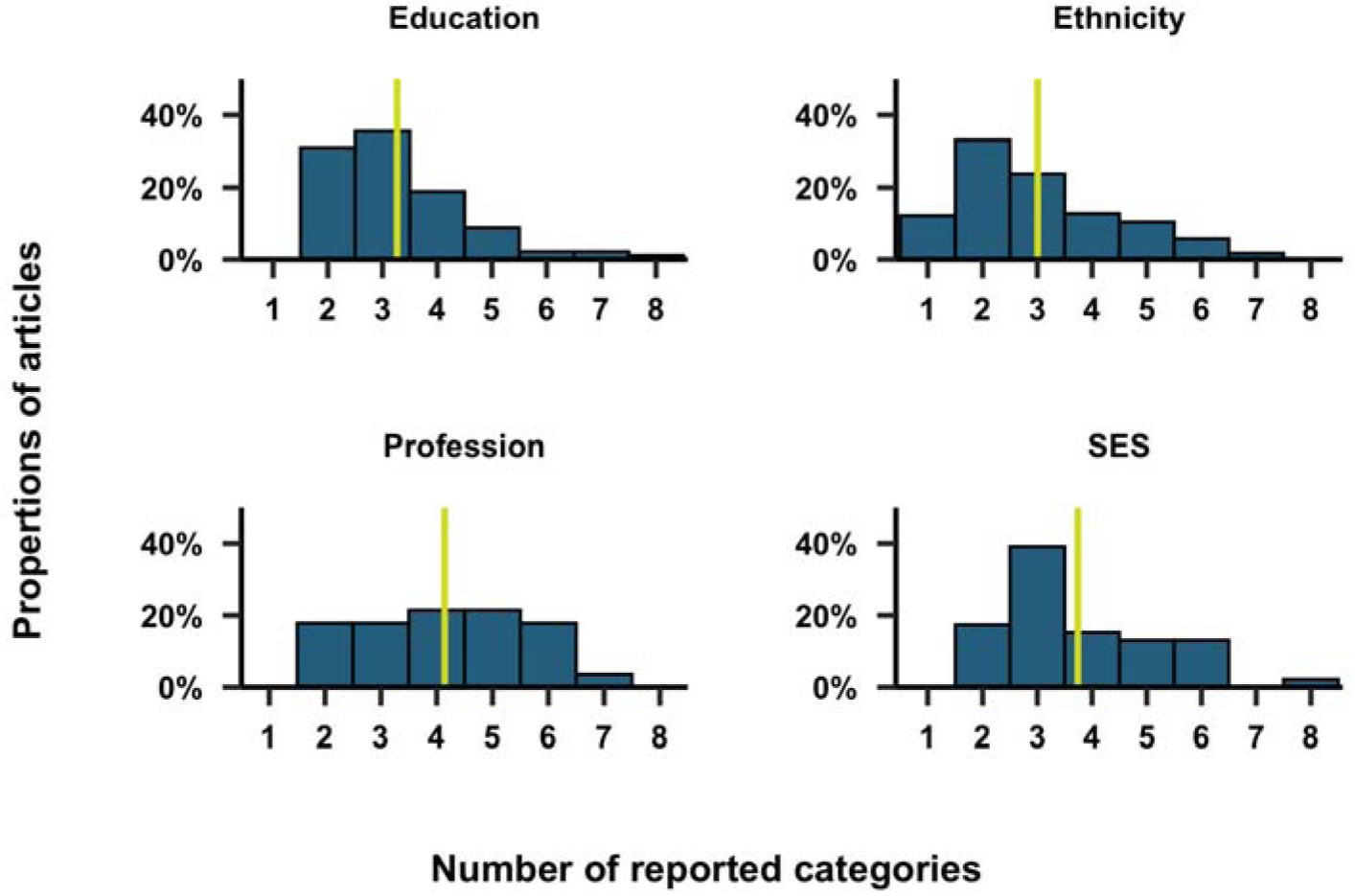
Distribution of the number of categories reported for education, ethnicity, profession and socio-economic status (SES). The yellow vertical line corresponds to the median.

### Study focus

We considered whether a given article reported on a specific, pre-defined group of people. We found that 3% of articles focused on a sex dependent feature, while 50% investigated a clinical feature. 1% of studies focused on twins, 1% on pregnant women, 2% on shift workers and 4% on university students.

### Analysis disaggregation

Sleep and circadian rhythms can be influenced by a variety of factors, including age, sex and ethnicity among others. By disaggregating data by these factors, researchers can identify patterns and associations that may not be apparent when analyzing the data as a whole. We thus investigated the extent to which articles reported subgroup analyses of the data based on one or more of the reported demographic variables. Over time (Fig. 11), we see a distinct evolution of the extent to which subgroup analyses of the study sample were performed. The most common subgroup analyses involve disaggregating by sex, age, or both.

**Figure 11.**
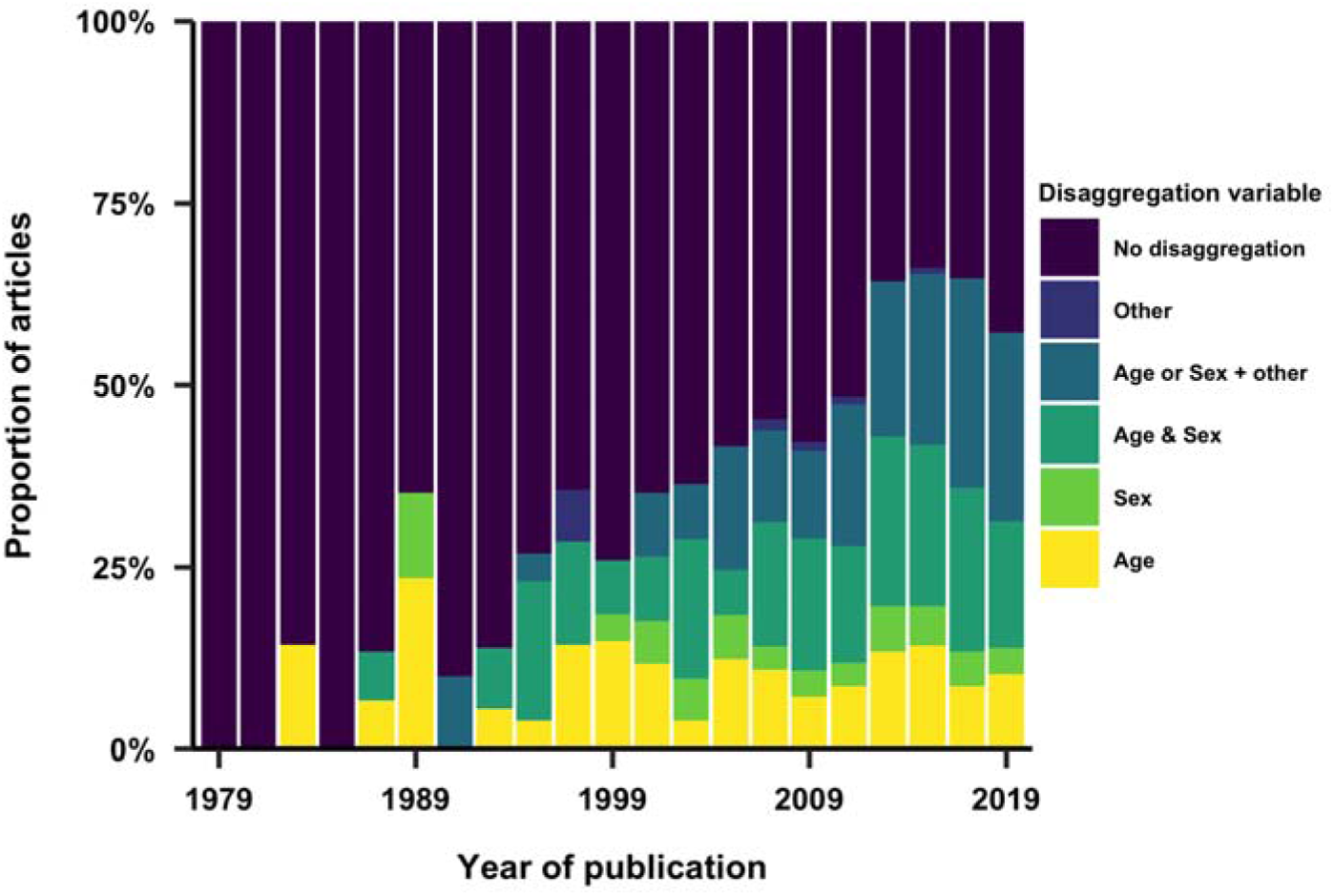
Use of study population characteristics as variables in the analysis.

## Discussion

### Summary of main findings

Here, we considered the inclusion and reporting of 1152 articles in chronobiology and sleep research, sampled from eight journals. Articles from recent years were more widely included, with articles originating in the US representing the largest group. While studies conducted in in North America and central Europe were predominant, those in Africa only represented 0.3% of all screened articles, underlining a gap in geographical inclusion. Over time, we found an increase in the range of sample sizes, as well as a wider spread of average ages included in the studies. We also observed an increase in the reporting of demographic variables (ethnicity, education, profession and socioeconomic status), with ethnicity being the most-reported variable in recent years. A wide variety of reporting strategies, which involve both objective and subjective measures, was also observed.

### Taking an inventory of represented study samples reveals the representativeness of our collective knowledge

The ability to generalize findings from the scientific literature to wide and diverse populations hinges upon the representativeness of the study sample with respect to demographic categories. The question to what extent the composition of a given study sample can make the generalizability of findings difficult or impossible has received attention in the field of psychology, where many articles published in prominent journals reflected participants from WEIRD (Western, Educated, Industrialized, Rich, and Democratic) contexts (61,62). In other fields, analyzes similar to the one in the present review have been published (42,63–65), but to our knowledge, this review represents a first look at the inclusion, reporting and analysis of participant demographics in chronobiology and sleep research.

### The need to consider individual differences

There is convergent knowledge that health and sleep and circadian physiology are subject to large individual differences and that a one-size-fits-all approach to promoting healthy sleep and circadian rhythms may not be effective. Demographic variables provide a lens through which to understand these individual differences, and importantly can provide insights into systemic disadvantages and inequities. In the clinical domain, the need to time therapy based on a patient’s individual circadian rhythm has more recently become the focus of the emerging field of chronotherapy or chronotherapeutics (66–69). Understanding interindividual variability needs to become a key research area to understand the circadian and sleep physiology in the face of human diversity.

### Advancing inclusion and diversity of study populations

Several efforts have been put in place to address gaps in inclusion and diversity in sleep and circadian rhythms research. Some funding agencies now require or strongly encourage researchers to report on demographic characteristics of study participants in grant applications and research publications. Many scientific journals also require authors to report on demographic characteristics of study participants, including age, sex, and race/ethnicity. Some journals also have specific policies to promote inclusion and diversity in research, such as requiring authors to explain any potential biases in their study design. Several initiatives, such as the the Sleep Research Society’s Diversity, Equity, and Inclusion Task Force are working to raise awareness of diversity issues in sleep research and develop best practices for promoting inclusion. Researchers are also increasingly working with community organizations to recruit study participants from underrepresented groups.

### Limitations of the current review

We turn to possible limitations of this review and the included analyses and discuss how they might introduce bias in our findings. First, we consider the possibility that the article selection procedure may have introduced biases. Our review only concerned articles from a subset of eight specialized journals written in the English language. As a consequence, the included articles were necessarily published in these journals, ignoring relevant articles published in other specialized journals (such as those included in the list of candidate journals), and articles published in other interdisciplinary journals. This raises the question to what extent we may have missed a section of the literature that would have been relevant to include in this study. As an alternative strategy, we considered randomly sampling a subset of chronobiology and sleep research articles produced by a general search (e.g. on search from “sleep OR chronobiology” on MEDLINE), but considered this to be too permissive. Our strategy of selecting a subset of candidate journals provided a reasonable trade-off, as well as sampling from a range of field-specific outlets.

Due to the non-uniform distribution of publication years of the included articles (Fig. 1), variables derived from published papers and visualized and/or analyzed by year will have varying uncertainty, with reported percentages from publications of the earlier years being most uncertain. The fact that early years are represented with fewer articles, however, is not a function of our data set, but of the exponential growth of scientific output (70–72).

### Towards standardized reporting of demographic variables: From checklists to schemas?

There are guidelines and/or checklists for standardizing reporting of participant characteristics, such as CONSORT (73) or STROBE (74) (an extensive database for health research reporting guidelines is provided by the Equator Network, https://www.equator-network.org/). Some biomedical journals (e.g. 75) specifically state demographic reporting requirements in the author instructions. Similarly, some organizations may make recommendations of specific reporting items for specific types of study questions (76). By following these guidelines and checklists, researchers can ensure that they report relevant and comprehensive information on participant characteristics. This can improve the reproducibility and transparency of research, facilitate data sharing and integration across different studies and datasets, and promote more equitable and inclusive sleep and circadian rhythms research.

Yet, these guidelines and/or checklists are largely focused on *what* is reported and not *how* it is reported. There is, *a priori*, however, no reason to not develop and use a standardized and machine-readable schema for reporting participant characteristics. The FAIR principles state that data should be findable, accessible, interoperable and reusable (77), and one way of realizing these criteria is the use of data schemas which could prescribe categories of data and common naming schemes for reporting participant characteristics. Importantly, however, “what gets counted counts” (78), and it will be imperative to understand to what extent such data schema may be exclusionary, e.g., by enforcing sex binaries (79), and whether any specific demographic variable is truly important (following the principle of data minimization). A further central point to consider is the extent to which disaggregation by demographic variables could be used to do harm in some way (78).

## Conclusion

This review provides a first look at the inclusion, reporting and analysis of demographic variables in the chronobiology and sleep research literature, considering >1000 articles across eight journals. We address the need to consider individual differences, as well as the dependence of sleep and circadian rhythms on demographic variables. Additionally, we outline an opportunity to improve reporting of participant-level characteristics using formalized data schemas.

